# Restructured mitochondrial-nuclear interaction in *Plasmodium falciparum* dormancy and persister survival after artemisinin exposure

**DOI:** 10.1101/2020.09.28.314435

**Authors:** Sean V. Connelly, Javier Manzella-Lapeira, Zoë C. Levine, Joseph Brzostowski, Ludmila Krymskaya, Rifat S. Rahman, Angela C. Ellis, Shuchi N. Amin, Juliana M. Sá, Thomas E. Wellems

**Affiliations:** Laboratory of Malaria and Vector Research, National Institute of Allergy and Infectious Diseases, National Institutes of Health Bethesda, Maryland, USA; Laboratory of Immunogenetics, National Institute of Allergy and Infectious Diseases, National Institutes of Health, Bethesda, Maryland, USA

**Keywords:** malaria, artemisinin-based combination therapy, drug resistance, AiryScan microscopy, fluorescence lifetime imaging, mitochondrial retrograde response

## Abstract

Artemisinin and its semi-synthetic derivatives (ART) are fast acting, potent antimalarials; however, their use in malaria treatment is frequently confounded by recrudescences from bloodstream *Plasmodium* parasites that enter into and later reactivate from a dormant persister state. Here we provide evidence that the mitochondria of dihydroartemisinin (DHA)-exposed persisters are dramatically altered and enlarged relative to the mitochondria of young, actively replicating ring forms. Restructured mitochondrial-nuclear associations and an altered metabolic state are consistent with stress from reactive oxygen species. New contacts between the mitochondria and nuclei may support communication pathways of mitochondrial retrograde signaling, resulting in transcriptional changes in the nucleus as a survival response. Further characterization of the organelle communication and metabolic dependencies of persisters may suggest strategies to combat recrudescences of malaria after treatment.

**IMPORTANCE:** The major first-line treatment for malaria, especially the deadliest form caused by *Plasmodium falciparum*, is combination therapy with an artemisinin-based drug (ART) plus a partner drug to assure complete cure. Without an effective partner drug, ART administration alone can fail because of the ability of small populations of blood-stage malaria parasites to enter into a dormant state and survive repeated treatments for a week or more. Understanding the nature of parasites in dormancy (persisters), and their ability to wake and reestablish actively-propagating parasitemias (recrudesce) after ART exposure, may suggest strategies to improve treatment outcomes and counter the threats posed by parasites that develop resistance to partner drugs. Here we show that persisters have dramatically altered mitochondria and mitochondrial-nuclear interactions associated with features of metabolic quiescence. Restructured associations between the mitochondria and nuclei may support signaling pathways that enable the ART survival responses of dormancy.

---

Artemisinin provides the basis for first line antimalarial treatment worldwide, particularly against the deadliest form of malaria caused by *Plasmodium falciparum* (1). Artemisinin and its semi-synthetic derivatives (collectively abbreviated here as ART) are among the best antimalarials for rapid parasitemia clearance and resolution of illness (2, 3). Yet, since its introduction, frequent recrudescences have been reported after ART monotherapy, necessitating the use of partner drugs for their prevention (4). Dormant intraerythrocytic parasites (persisters) have been found to produce these recrudescences (5-8), but much remains to be understood about the nature of these persisters and how they develop from only a small fraction (less than ~1% (5, 7)) of ART-treated populations.

Although dormancy of intrahepatic parasites (hypnozoites) of certain *Plasmodium* spp. was well known, it was not until 1995 that small numbers of dormant intraerythrocytic parasites were reported from *P. falciparum* populations treated with pyrimethamine or with sequential passages through a D-sorbitol solution to destroy actively replicating parasites (9). Importantly, the dormant parasites were able to survive these treatments for several days, after which they gave rise to recrudescent populations that were as susceptible to pyrimethamine or D-sorbitol as the original populations. Further, no relationship could be demonstrated between the incidence or timing of recrudescence and drug concentration (10).

Dormant forms following ART exposure have been distinguished from actively replicating ring stages and pyknotic parasites in Giemsa-stained thin blood films by their small rounded appearance with magenta colored chromatin and condensed blue cytoplasm (6, 8). Outgrowth experiments using Rhodamine 123 staining demonstrated that mitochondrial membrane potential is critical to persister viability (11). Enzymes involved in fatty acid biosynthesis, pyruvate metabolism, and the isoprenoid pathway are among the few upregulated pathways in dormant parasites following ART exposure (12, 13), while expression changes of cyclin and cyclin dependent kinase (CDK) genes have been correlated with dormancy and reactivation of persister forms (14).

Considerably different *in vitro* recrudescence times can be found with *P. falciparum* parasites that have distinct genetic backgrounds, e.g. between the 967R539 and KH002-009 lines studied by Breglio et al. (15). However, dependence of recrudescence time on *P. falciparum* K13 protein (PfK13) polymorphisms could not be demonstrated, as the 967R539 and KH002-009 lines were both PfK13 wild-type; further, recrudescences times were not significantly faster in comparative evaluations of PfK13 R539T- or C580Y-*vs*. wild-type alleles in pairs of isogenic clones (15, 16). These findings contrasted with the effects of PfK13 mutations on the parasite survival in Ring Stage Assays (RSAs), wherein early ring-stages (0–3 h post invasion) are exposed for 6 hours to a physiological level of ART (16, 17). However, because the RSA phenotype does not include the PfK13-independent susceptibilities of more mature trophozoite- and schizont-stages, it does not correlate with standard half-maximum inhibitory concentrations (IC^50^) or clinical treatments that provide continual ART exposures for more than one intraerythrocytic cycle (18). PfK13 status and the ring-stage phenotype are thus divorced from the ability of parasites to become dormant and survive ART monotherapy for periods of up to 10 days (19).

Recent advances in methods for fluorescence cell sorting, subcellular organelle imaging, and metabolic activity analysis offer new approaches to study persister forms. AiryScan Microscopy (ASM) provides a markedly improved resolution and signal-to-noise ratio relative to standard confocal microscopy for organelle-level imaging of individual cells (20). Fluorescence lifetime imaging (FLIM) with exogenous or endogenous (auto-fluorescent) fluorophores now enables non-invasive characterization of metabolic changes and provides insight into the redox status of tissues (21-23). Phasor analysis of FLIM data provides a convenient 2D graphical approach that has been applied to the metabolic characterization and responses of germ cells, bacteria, and keratinocyte cells (24-27).

Here we compare the subcellular structure and metabolic phenotypes of persisters to those of actively replicating ring forms from two clonal *P. falciparum* lines: GB4, an African line of Ghanaian origin, and 803, a Southeast Asian line from Cambodia. In previous work with parasites from a GB4×803 cross, a standard 3-day course of artesunate was shown to clear infections of non-human primates to microscopically undetectable levels, just as they do in humans, but frequent *in vivo* recrudescences occurred whether or not the parasites carried a PfK13 C580Y mutation from the Cambodian 803 parent (16). To investigate the events of dormancy that underlie these recrudescences, we have now used ASM to study the subcellular features of GB4 and 803 parasites, including changes in the mitochondria and their proximity to nuclei that suggest mito-nuclear interactions. Autofluorescence FLIM-phasor analysis indicates increased free NADH in persisters relative to actively replicating ring forms, which is consistent with metabolic quiescence in dormancy.

## RESULTS

### Recrudescence after ART exposure is innate to *P. falciparum*

To further expand on previous observations that the recrudescence profiles of ART-treated *P. falciparum* populations can vary with their genetic background (7, 9, 14-16, 28), we followed cultures of synchronized GB4 and 803 parasites after their exposure as young rings to 700 nM DHA for 6 hours and three successive 5% D-Sorbitol treatments at 24, 48 and 72 hours. Fig. 1A shows the recrudescence curves, from the initial 2% ring parasitemias treated with the DHA, through the low levels (<0.05%) of parasites in dormancy, to the recovery of ~2% actively growing GB4 and 803 parasites at 20 and 16 days, respectively. Data from RSAs were also compared and showed lower percentage survivals of GB4 relative to 803 parasites, consistent with their difference by PfK13 C580Y and its effect in the GB4×803 cross (16) (see Fig. S1A in the supplemental material). Taken together, these results along with the observations of previous studies reinforce that recrudescences after ART treatment are universal features of *P. falciparum* lines and that variations of recrudescence time may appear to partner with PfC580Y in some cases, and in other cases not, with the effects of different genetic backgrounds.

**FIG 1.**
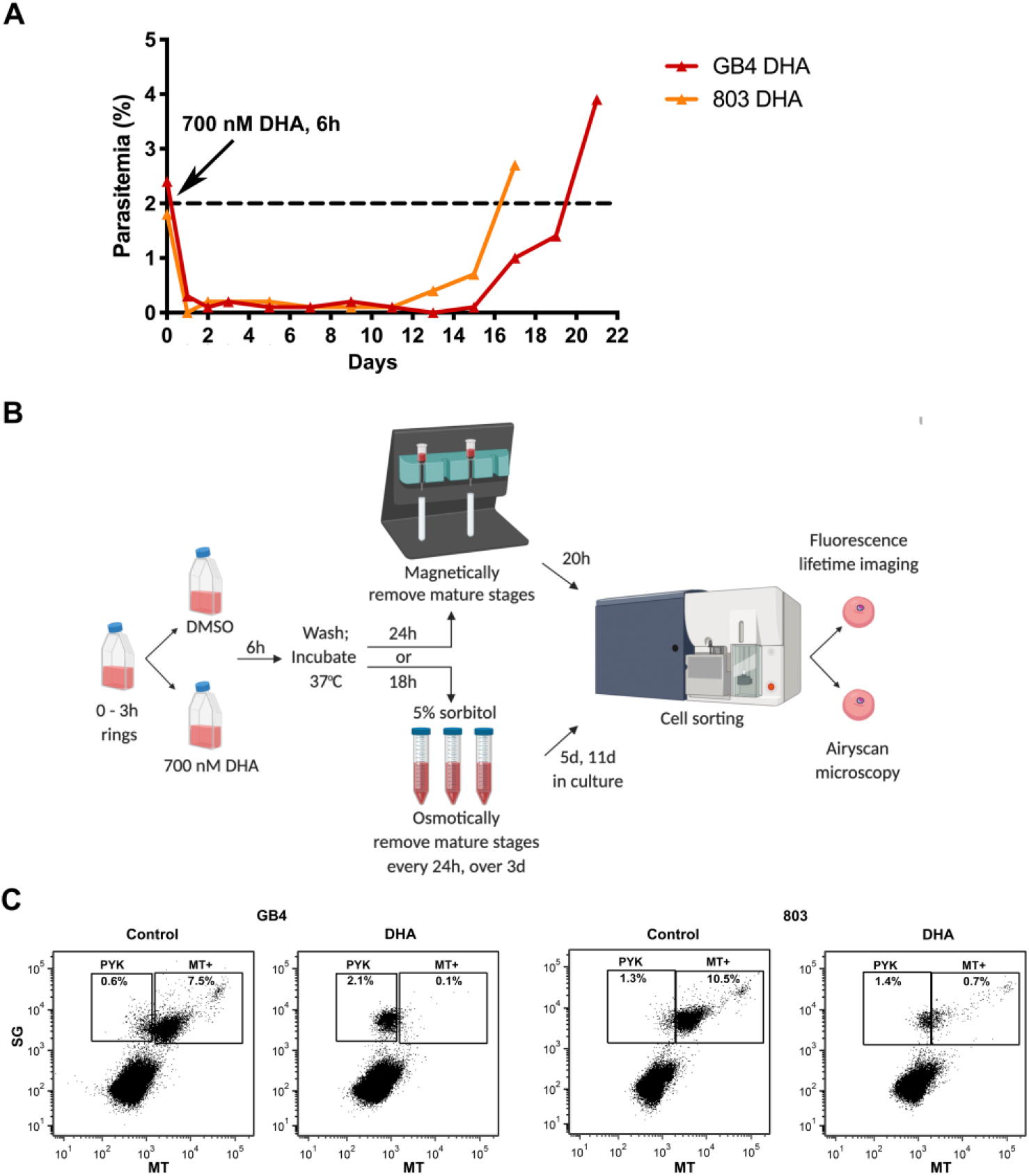
Recrudescence of DHA-treated *P. falciparum* parasites *in* vitro and sorting of their dormant persister forms by FACS. (**A**) Recrudescence curves of GB4 and 803 parasites after exposure of early ring stages to 700 nM DHA for 6 hours and three daily 5% D-sorbitol treatments. (**B**) Schematic of GB4 and 803 parasite preparations for study by ASM and FLIM-phasor analysis. The parasites were synchronized through two sorbitol treatments, 46 hours apart, to obtain 0 – 3 hour rings. At the start of the experiment (*t* = 0 hours), the GB4 or 803 populations of 0 – 3 hour rings were treated at 2% parasitemia with either 700 nM DHA or 0.1% DMSO vehicle for 6 hours, washed and returned to culture. In one arm of the study, DHA treated parasites at *t* = 30 hours were passed through a magnetic depletion column to remove mature-stage parasites that had continued to grow after DHA treatment. The recovered parasites were then returned to culture on a rocking incubator for 20 hours at 37^°^C along with the vehicle control parasites. At *t* = 50 hours, 1mL of each culture was stained with either SG plus MT for FACS and ASM, or with MT only for FACS and FLIM-phasor autofluorescence analysis. In the other arm of the study, parasite populations exposed to DHA or DMSO only were given three daily treatments with 5% sorbitol to select for persister forms. On days 5 and 11, 1 mL of each culture was stained and studied in the same way as the magnetically-purified by ASM and FLIM-phasor autofluorescence analysis. (**C**) SG-positive parasites at *t* = 50 hours were gated into two populations based on MT fluorescence intensity. Large numbers of pyknotic forms in the PYK gates of the DHA treated populations represent parasites killed by the DHA. Parasites from the MT+ gate were used for ASM. SG = SYBR Green I, MT = MitoTracker Deep Red FM. Uninfected erythrocytes that lack SG and MT signals are indicated in the lower left quadrant of the plots.

### Viable persister forms can be sorted by a fluorescence sensor of mitochondrial potential

To identify and isolate persister parasites after DHA treatment of synchronized ring stages, we adapted the method of Peatey et al. (11) to use MitoTracker Deep Red FM (MT) instead of Rhodamine 123 as a sensor of mitochondrial membrane potential. The DHA-treated samples were passed through a magnetically-activated cell sorting (MACS) column at *t* = 30 hours to deplete mature-stage, hemozoin-containing parasites, returned to culture, and finally stained with MT and SYBR Green I (SG, for nucleic acid staining) before Fluorescence Activated Cell Sorting (FACS) at *t* = 50 hours (Fig. 1B).

FACS dot plots of the sorted GB4 and 803 samples are shown in Fig. 1C. In the upper left quadrant of each plot, the gate region labeled “PYK” includes pyknotic parasites that lacked mitochondrial potential whereas, in the upper right quadrant of the plot, the gate region labeled MT+ includes parasites that were positive by MT for mitochondrial activity. In this MT+ gate region, the SG and MT signals that extend upward and to the right indicate the progression of young ring stages to mature stage parasites, as described with fluorescent markers in previous studies (29). FACS patterns of the control vehicle-treated GB4 or 803 populations in this MT+ region (Fig. 1C, first and third panels) also show a dense cluster at the lower left and a lighter cluster at the upper right which, considering the timeline of development and reinvasion in the control populations, can be explained by the number of ring stages at the beginning of the second intraerythrocytic cycle relative to fewer schizonts that have not yet finished the end of the first cycle (see Fig. S1B in the supplemental material). In contrast, FACS results from the DHA-treated parasites (Fig. 1C, second and fourth panels) registered large numbers of parasites that lack mitochondrial potential in the PYK region, as expected after pyknosis and cell death from the drug. To the right of the PYK region, viable GB4 and 803 parasites registered in the MT+ region where fewer GB4 than 803 counts agree with their different RSA survival levels (see Fig. S1A in the supplemental material).

Peatey et al. (11) showed that parasites that had been DHA-treated and passed through a magnetic column at *t* = 30 hours were enriched for dormant persister forms. Since the enrichment is imperfect, FACS at *t* = 50 hours would be expected to identify persisters along with some actively replicating mature stages that escape removal by the single magnetic column. We tested this expectation by counting the parasite stages in Giemsa-stained thin blood films from the culture samples immediately before FACS. In three independent experiments using parasite populations that had been passed through the magnetic column at *t* = 30 hours, blood films at *t* = 50 hours showed that persisters in the control vehicle-treated populations were greatly outnumbered by rings, whereas overall greater fractions of persisters and pyknotic forms were present in the DHA-treated populations (see Fig. S2 in the supplemental material). Although the occurrence of ring-stage parasites in these DHA-treated populations were relatively infrequent, the median and interquartile range (IQR) of the GB4 ring stage percentages (0.90 (0.25 – 1.79) %) was less than that of 803 ring stages (8.04 (7.13 – 15.73)%; p = 0.028), consistent with the lower survival of GB4 relative to 803 parasites in RSA determinations.

Outgrowth experiments were performed to confirm the viability of the sorted parasite populations. For this purpose, cultures were seeded with parasitized erythrocytes that had been exposed as young ring forms to vehicle control or DHA, the DHA treated sample was magnetically purified at *t* = 30 hours, then both samples were sorted and collected at *t* = 50 hours by a gating strategy designed to separate SYBR green positive cells having the dim MT fluorescence of pyknotic forms (SG/MT^D^), the intermediate MT fluorescence of persisters and ring stages (SG/MT^I^), or the bright MT fluorescence of schizonts (SG/MT^B^) (see Fig. S3 in the supplemental material). Giemsa-stained thin films of these FACS collections showed that the collections preponderantly contained pyknotic cells, persisters, or mature stages, respectively, although the amounts of material were insufficient to reliably count the small fractions of admixed stages (see Fig. S4 in the supplemental material).

Days to 0.5% parasitemia outgrowth were determined for these different populations in two replicate experiments with GB4 parasites and one experiment with 803 parasites. Results showed outgrowth in 14 – 20 days from wells seeded with 30,000 of the collected DHA-treated SG/MT^I^ GB4 or 803 parasites (see Tables S1 – S3 in the supplemental material). As described above and documented in Fig. S2, the majority of the SG/MT^I^ seeded parasites were persisters, although a minority of active rings were present (fewer from GB4 than from 803) which also could have contributed to the outgrowth. Outgrowth was also observed at 17 – 27 days from the collected SG/MT^B^ mature stage 803 and GB4 parasites seeded at 1,200 – 6, 000 parasites/well. In contrast, except for a single late instance of recovery at 28 days, no outgrowth was obtained from the pyknotic SG/MT^D^ forms collected after DHA treatment, supporting the utility of MT as an indicator of viability for parasites exposed to the drug.

### AiryScan Imaging Reveals Organelle Level Changes in GB4 and 803 Persisters

Isolated persister and ring-stage forms were imaged by high-resolution ASM and compared for fluorescent mitochondrial and nuclear volumes, the shapes of these volumes as indicators of organellar morphology, and the mean distances between these volumes as indicators of relative mito-nuclear positioning.

Fig. 2 shows representative surface renderings of the volumes of nuclear DNA and mitochondrial fluorescence regions from experiments with vehicle control- or DHA-treated GB4 and 803 parasites. In the control, the nuclear DNA (green) and mitochondrial (red) volumes of the ring forms were typically well separated from one another. Nuclear DNA volumes presented with an oblate or boxy appearance and were greater than the MT-stained volumes of mitochondria. The mitochondrial volumes were usually oblate in appearance, reminiscent of the rounded or elongated dot-like mitochondrial images reported from ring-stage parasites by Scarpelli et al. (30). Longer rod-like appearances of ring-stage mitochondria reported by van Dooren et al. (31) were not characteristic of the MT-stained mitochondria in our ASM images.

**FIG 2.**
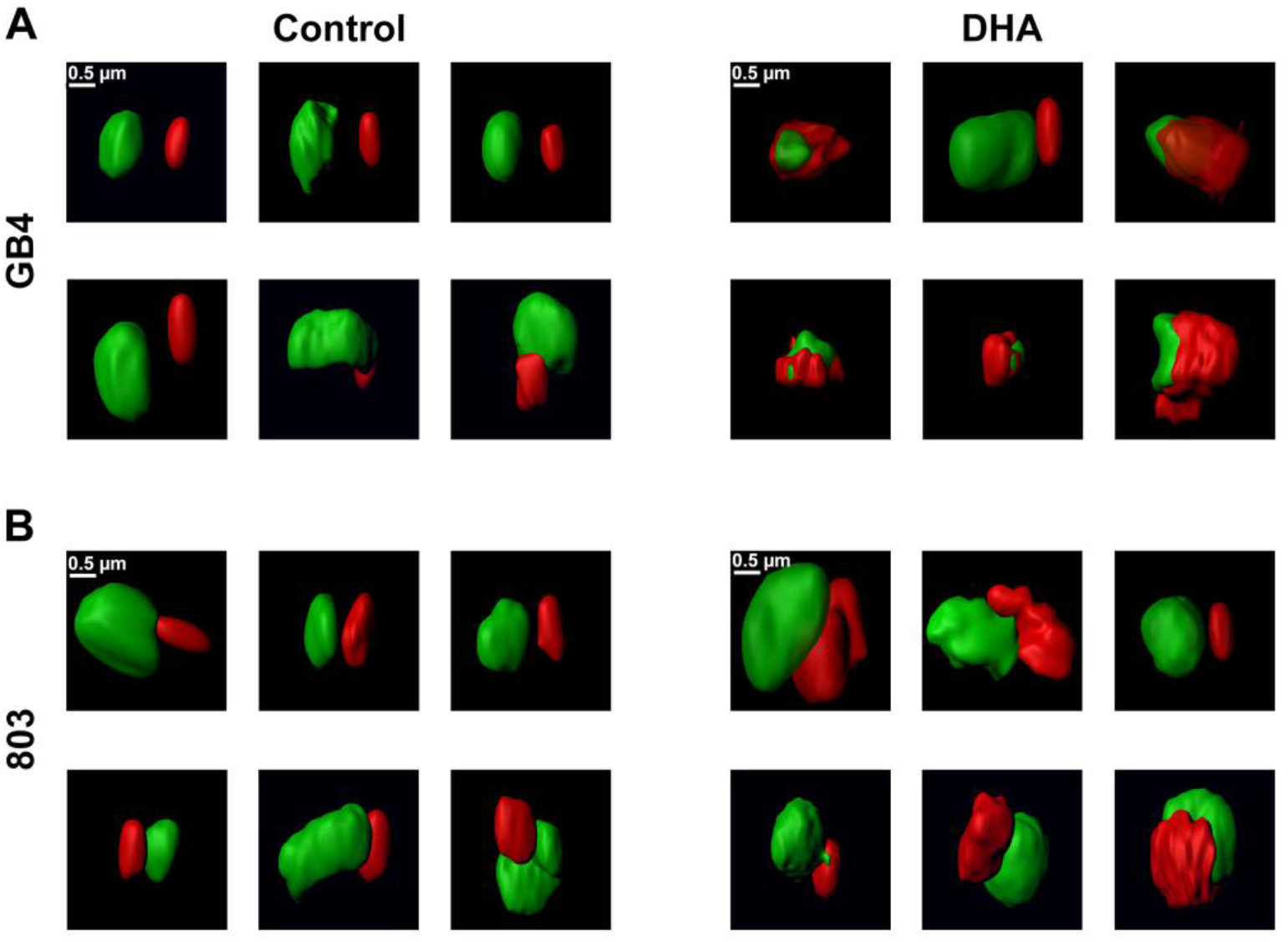
ASM images of control and DHA-treated GB4 and 803 parasites at *t* = 50 hours. Individual panels show processed ASM images of the mitochondrial (red) and nuclear (green) fluorescence signals from six representative parasites in each treatment group separated by FACS. (**A**) GB4 parasites from control 0.1% DMSO vehicle-treated samples present smooth, oblate mitochondrial and nuclear volumes. Images in this control group are consistent with young ring-stage parasites in the culture at *t* = 50 hours. DHA-treated parasites show the distinctly different morphologies of persisters, with rumpled and corrugated mitochondrial volumes that are in close approximation to the nuclei. (**B**) Images of the control 803 parasites at *t* = 50 hours also have the smooth, oblate appearance of mitochondria separated from the nucleus, as expected of actively replicating young parasite ring forms. Images of DHA-treated 803 parasites, in comparison to those of the treated GB4 population, are consistent with a mixed population of cells: many with the rumpled, corrugated mitochondria characteristic of persisters, while others have the smooth, oblate mitochondria apart from nuclei that is characteristic of ring-stage parasites.

After DHA treatment of synchronized GB4 parasites followed by magnetic column separation at *t* = 30 hours, actively growing parasites were mostly eliminated, and surviving persisters greatly predominated in the DHA-treated GB4 population captured at *t* = 50 hours (Fig. 1C second panel, and Fig. S2 in the supplemental material). Fig. 2A (right panel) presents the nuclear and mitochondrial morphologies of six parasites from this population: one shows a smooth oblate mitochondrion separate from the nucleus that is characteristic of an actively growing ring-stage form, whereas the five others exhibit distinctly different appearances, with rumpled and corrugated mitochondria in close approximation to the nuclei. The mitochondria partially enwrap the nuclei, and many are enlarged compared to the control ring-stage parasites. Contact sites appear to be present between the nuclear DNA and mitochondria as another potential feature of the persister phenotype.

Because single column separation at *t* = 30 hours does not completely remove actively replicating parasites from the DHA-treated 803 parasites (Fig. 1C, fourth panel, and Fig. S2 in the supplemental material; daily column separations would have been required for more complete purification of the persisters (7)), the parasites imaged by ASM at *t* = 50 hours comprised a major population of persisters with an admixture of ring-stage forms. Thus, in comparison to the DHA-treated GB4 population, images from the DHA-treated 803 population show more frequent examples of cells with the smooth surfaced, normal volume mitochondria separate from the nuclei, as observed in the control 803 population (Fig. 2B, right panel). Other images of DHA-treated 803 parasites have rumpled, corrugated mitochondria in close approximation to nuclei as observed in the DHA-treated GB4 persisters (Fig. 2B, right panel). These findings confirm the presence at *t* = 50 hours of a mixed population of persisters and actively replicating ring stages in the DHA-treated 803 sample, consistent with the results shown in Fig. 1C and Fig. S2.

### Population Measures of Mito-nuclear Distance and Organelle Volumes Differ between Persisters and Actively Replicating Ring Stages

To quantify the phenotypes of persisters *vs*. actively replicating ring-stage parasites, mito-nuclear distances, mitochondrial volumes, and nuclear volumes were determined from parasites of the control vehicle- and DHA-treated GB4 and 803 populations. At *t* = 50 hours, a mito-nuclear median distance and IQR of 0.56 (0.32 – 0.98) µm was obtained from the control, untreated population of GB4 ring-stages (*n* = 167), whereas a much smaller median distance of 0.19 (0.11 – 0.28) µm was obtained from the population of DHA-treated GB4 parasites that predominantly consisted of persisters (*n* = 167) (Fig. 3A; p < 0.0001, unpaired t-test). In the case of 803, mito-nuclear median distance and IQR in control vehicle-treated population of ring-stages was 0.39 (0.25 – 0.53) µm (*n* = 190), not significantly smaller than 0.31 (0.17 – 0.49) µm (*n* = 158) obtained for DHA-treated 803 parasites (Fig. 3A; p = 0.50, unpaired t-test). The insignificance of this difference between the control and DHA-treated 803 populations is consistent with a greater presence of surviving active ring-stages with persisters in the DHA-treated 803 population.

**FIG 3.**
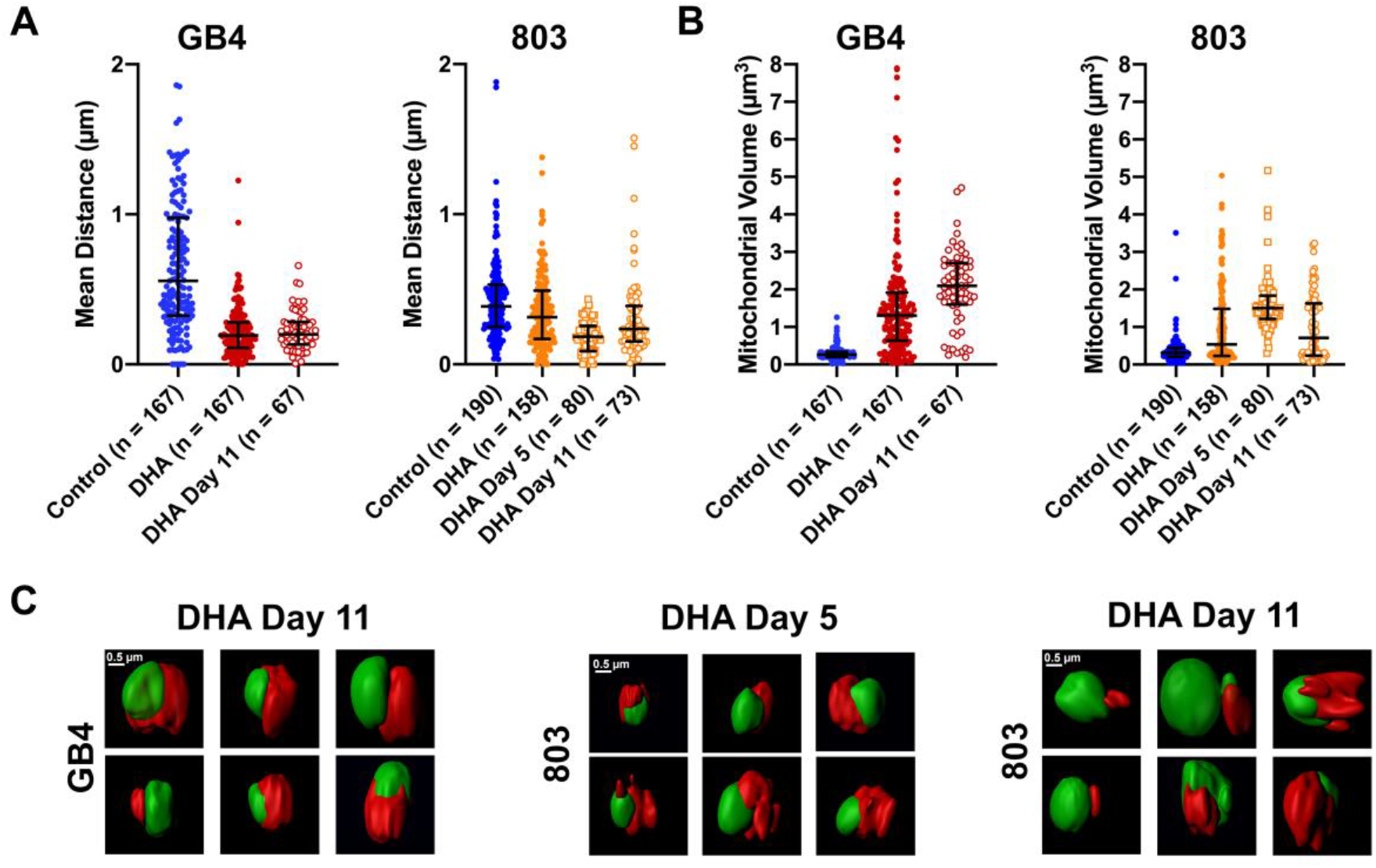
Quantifications of mito-nuclear distances and mitochondrial volumes of GB4 and 803 parasites at *t* = 50 hours, Day 5, and Day 11 after DHA treatment. (**A**) Scatterplots, median, and interquartile ranges (IQR) of mito-nuclear distances in the populations of GB4 and 803 parasites exposed to 0.1% DMSO (control) or 700 nM DHA for 6 hours, selected, sorted, and analyzed as described in the text. In the mito-nuclear distance scatterplots, there were five outliers beyond the bounds of the y-axis in the GB4 Control group (at 2.8 µm, 2.5 µm, 2.2 µm, 2.7 µm, and 6.8 µm), one outlier in the 803 Control group (at 3.7 µm), and three outliers in the 803 DHA group (at 2.6 µm, 2.0 µm, 5.5 µm). (**B**) Mitochondrial volume scatterplots, median, and IQR from the same populations used for mito-nuclear distance determinations. (**C**) Middle panel presents representative ASM images of DHA-treated 803 parasites on Day 5 post treatment. The six examples of persisters show rumpled, corrugated mitochondria in close approximation to the nuclei. The right and left panels present the surface images of DHA-treated GB4 and 803 parasites at Day 11. Persister morphologies are evident in the images from each parasite line, including rumpled and corrugated mitochondria in close approximation to the nuclei.

Mitochondrial volume comparisons of control and DHA-exposed GB4 or 803 parasites revealed an increase in the mitochondrial volume 50 hours after DHA treatment. In control GB4 parasites the median mitochondrial volume and IQR was 0.26 (0.20 – 0.34) µm^3^, whereas the post DHA treatment mitochondrial median volume was 1.30 (0.64 – 1.9) µm^3^, a 5-fold increase (Fig. 3B; p < 0.0001, unpaired t-test). In control 803 parasites, the median mitochondrial volume was 0.31 (0.20 – 0.43) µm^3^ and the DHA-exposed median volume was 0.54 (0.23 – 1.5) µm^3^ (Fig. 3B; p < 0.0001, unpaired t-test). Compared to the results from GB4 parasites, the less dramatic increase of median mitochondrial volume in DHA-exposed 803 parasites at *t* = 50 hours may be partially explained by its mixed population of persisters, with large mitochondrial volumes, plus actively replicating ring forms, with smaller mitochondrial volumes. We note that mixed populations of actively replicating and persister forms have been detected in post-artesunate blood smears of patients infected with parasites carrying the PfK13 Kelch propeller R539T mutation but not with parasites carrying wild-type PfK13 (28). These findings with GB4 and 803 parasites are consistent with the increased mitochondrial volumes that have been reported from other studies of ART-treated *Plasmodium* and *Toxoplasma gondii* (32, 33).

### Isolation and Characterization of Persister Parasites 5 and 11 Days after DHA Treatment

After examining the morphological characteristics of persister mitochondria and quantifying their close apposition to nuclei at *t* = 50 hours, we looked for the presence of these features in persisters 1 – 2 weeks old, before their recrudescence in culture. Samples of the DHA-treated parasite populations were examined and counted by Giemsa-stained thin smear microscopy at Day 5 and 11, sorted by MT intensity in FACS, then collected and subjected to ASM and analysis (Fig. 1B).

Counts of the Giemsa-stained samples confirmed the presence of persisters among pyknotic forms in the DHA-treated GB4 and 803 populations at both days 5 and 11. Low percentages of actively replicating parasites were also observed, more frequently on day 11 than day 5, particularly in the 803 population where these active forms accounted for approximately 6% of the total count relative to 19% persisters and 75% pyknotic forms (see Fig. S2 in the supplemental material). This increased count of 803 active forms on day 11 agrees with the earlier recrudescence of 803 relative to GB4 parasites two weeks after treatment with DHA (Fig. 1A).

ASM of the DHA-treated 803 parasites collected on day 5 showed nearly complete preponderance of persister morphology in which large and rumpled mitochondrial volumes often appeared to partially enwrap the nucleus (Fig. 3C, center panel). The collected GB4 parasites were not imaged on day 5 but on day 11, when features similar to those of persisters at *t* = 50 hours were present in virtually all of the cells (Fig. 3C, left panel). The collected parasites from the 11^th^ day 803 population also showed features of persisters in the majority of cells, but there were also many examples of cells with smaller oblate mitochondria separate from the nucleus (Fig. 3C, right panel; compare with the images in Fig. 2B). This evidence for actively growing ring-stage forms among persisters in the 11^th^ day 803 population is consistent with the observations from Giemsa-stained thin films and waking from dormancy at the beginning of the recrudescence curve shown in Fig. 1A.

In agreement with the nearly complete preponderance of persisters in the DHA-treated population on Day 5, the median mito-nuclear distance of the 803 parasites on this day was at its smallest (0.18 (0.087 – 0.26) µm; Fig. 3A) while the median mitochondrial volume was at its largest (1.5 (1.2 – 1.8) µm^3^; Fig. 3B). By Day 11, the median mito-nuclear distance of the 803 population was again larger (0.24 (0.15 – 0.39) µm) and the mitochondrial volume smaller (0.71 (0.24–1.6) µm^3^) than on Day 5, as expected from the presence of active stages that had waken from the persisters. The median mito-nuclear distance of the 803 population on Day 11 was also greater than that of GB4 (0.24 (0.15 – 0.39) µm *vs*. 0.20 (0.13 – 0.28) µm; p = 0.0187, unpaired t-test; Fig. 3A), and the median mitochondrial volume was comparatively smaller ((0.71 (0.24 – 1.6) µm^3^ vs. 2.1 (1.6 – 2.7) µm^3^; p < 0.0001; Fig. 3B), reflecting the greater fraction of actively replicating parasites in the 803 *vs*. GB4 population (Fig. S2).

Nuclear DNA volumes of the DMSO vehicle- and DHA-treated GB4 and 803 parasites were assessed at *t* = 50 hours and compared volumes of the DHA-treated populations at Days 5 and 11 (see Fig. S5 in the supplemental material). At *t =* 50 hours, the median nuclear DNA volume of DHA-treated GB4 parasites was smaller than that of vehicle-treated controls (0.83 (0.57 – 1.1) µm^3^ *vs*. 1.1 (0.85 – 1.4) µm^3^, p = 0.0012, unpaired t-test); however, the median volume of DHA-treated 803 parasites was relatively larger (2.1 (1.5 – 2.8) µm^3^ *vs*. 1.8 (1.2–2.2) µm^3^; p < 0.0001). The minimum median DNA volume of the 803 population occurred on Day 5 (1.2 (1.1 – 1.5) µm^3^), when the fractional content of persisters was highest in the population. One Day 11, the median DNA volumes of the GB4 and 803 parasites were not significantly different (1.5 (1.1 – 1.9) µm^3^) *vs*. 1.6 (1.2–2.2) µm^3^; p = 0.09, unpaired t-test).

We considered whether the separation of the mitochondria and nuclei could have been reduced by shrinking of the overall available space in the parasite after drug treatment. One of our controls for preparation quality was to assess infected erythrocyte areas of the vehicle-treated and DHA-treated samples. Cells from FACS collections were attached onto Poly-L-lysine coated surfaces and imaged by differential interference contrast (DIC) microscopy before fluorescence lifetime imaging. Analysis of these images found no reduction in the median surface area of DHA-treated relative to vehicle only-treated cells; in fact, the data suggested a possibly larger surface area of the DHA-treated population (supplemental Fig. S6). Thus, with the caveat that the ability of the vehicle control- and DHA-treated cells to flatten may differ, these assessments provided no support for differential constriction of the cells due to DHA-treatment.

### FLIM Metabolic Imaging Distinguishes the States of Actively Replicating Parasites and Persisters

We employed FLIM imaging to assess ratios of the bound and free states of nicotinamide adenine dinucleotide (NADH) as an autofluorescence indicator of the cellular metabolic state (23, 25, 34, 35) (Fig. 4A). FLIM data were collected from actively replicating control and DHA-treated populations at *t* = 50 hours and subjected to phasor analysis in three independent experiments (Fig. 4B). Results from GB4 parasites showed that the phasor distribution of the DHA-treated population was shifted to the right of the untreated control ring stages, indicative of a greater level of free NADH and quiescent state of the persister phenotype (control: (0.75, 0.22), DHA-treated: (0.80, 0.19); p < 0.0001, Pillai trace). With the 803 parasites, the DHA-treated population had only a higher S coordinate than the control group (control: (0.63, 0.14), DHA-treated: (0.63, 0.16); p < 0.05, Pillai trace). Little or no shift of the phasor position to the right was consistent with the presence of metabolically active ring forms along with dormant persisters in the DHA-treated 803 population at *t* = 50 hours.

**FIG 4.**
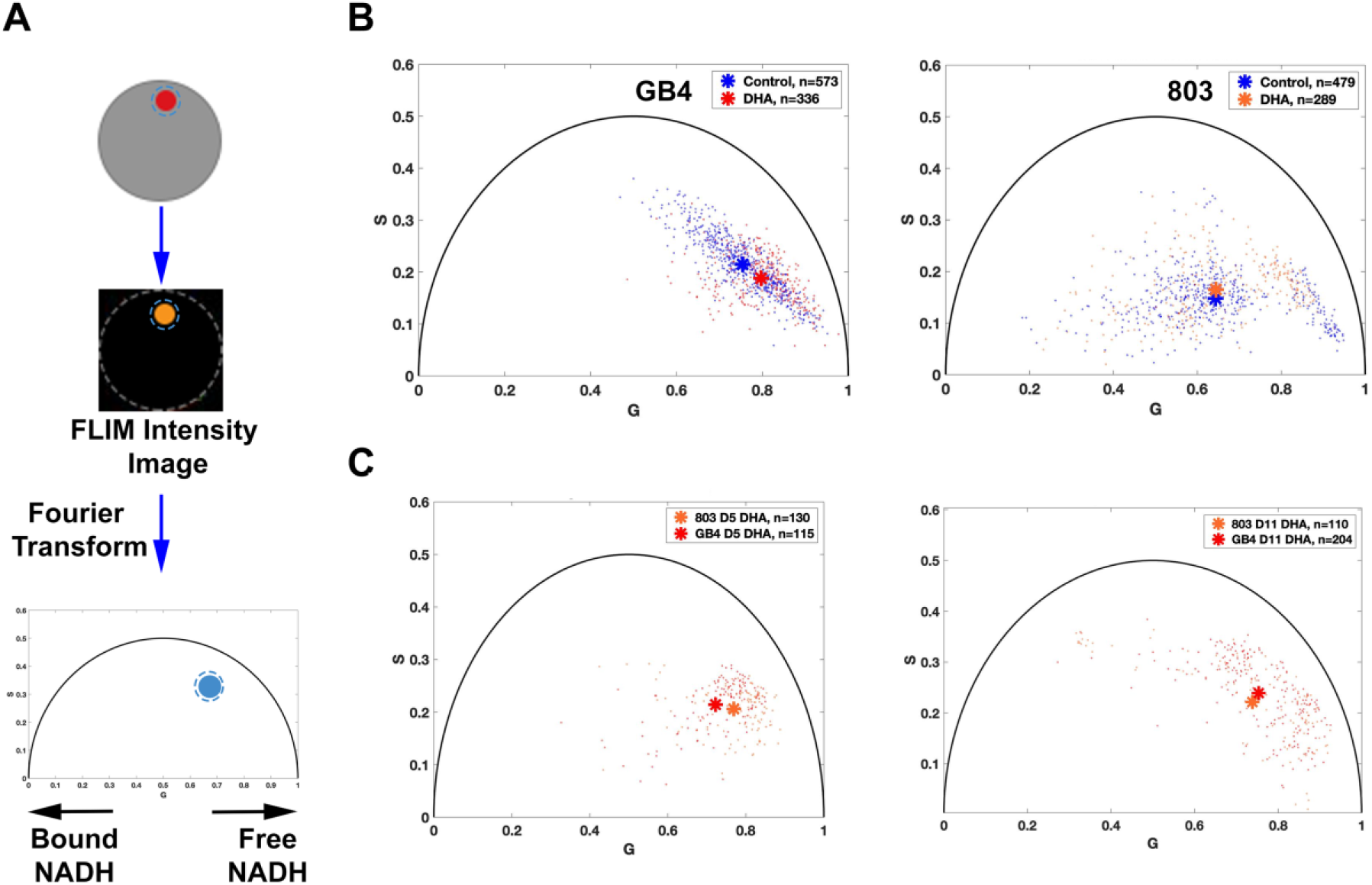
Metabolic phenotyping of untreated control *vs*. DHA-treated GB4 and 803 parasites. (**A**) Schematic flow of FLIM-phasor data collection and analysis. Parasites were stained only with MT, sorted by FACS, and seeded into a 10 × 10 µm polydimethylsiloxane (PDMS) stencil microwell covered with 0.01% poly-lysine. The region of interest (ROI) in a parasite is established by the boundary of MT fluorescence, and this ROI mask is applied to the lifetime datafile represented by its FLIM intensity image. At least ten million photon events are counted while limiting the time of exposure of these parasites to 750 nm excitation. Next, Fourier transformation of the FLIM data is performed on the data from each pixel and the frequency domain results are averaged to represent a data point on the 2D phasor plot. Each position has coordinates G (from 0 to 1) and S (from 0 to 0.5); single exponential decays fall upon the semicircle whereas complex multiexponential decays, such as occur with fluorescent lifetimes of metabolic coenzymes, fall within the semicircle. A shift to the right (shorter fluorescence lifetime) indicates increased free NADH whereas a shift to the left (longer lifetime) indicates increased enzyme-bound NADH. (**B**) Autofluorescence FLIM-phasor graphs from the GB4 and 803 parasite populations at *t* = 50 hours. The phasor distribution of untreated control GB4 parasites has a mean coordinate of (0.7542, 0.2150) whereas the distribution of DHA-treated parasites has a mean coordinate of (0.7971, 0.1879) (p < 0.0001, Pillai trace). This shift after DHA treatment is toward increased free NADH, consistent with the quiescent metabolic state of persisters. The distribution of untreated 803 parasites at *t* = 50 hours (right) has a mean coordinate of (0.6341, 0.1400) and the phasor distribution of DHA-treated parasites has a mean coordinate of (0.6270, 0.1570). The distribution of DHA-treated 803 parasites is slightly shifted upward relative to control (p < 0.05, Pillai trace), but not to the left or right, consistent with the presence of actively replicating 803 parasites that survived ring-stage exposure to DHA. Asterisks mark the mean phasor positions of the control and DHA-treated parasite distributions. Comparison of intensity images in the same field of view before and after FLIM acquisition confirmed little or no photobleaching. Figure created using Biorender.com software. (**C**) Autofluorescence FLIM-phasor graphs for DHA-treated GB4 and 803 on Day 5 and Day 11. In the phasor distributions from Day 5 (left), DHA-exposed GB4 and 803 parasites have mean coordinates of (0.72, 0.21) and (0.77, 0.21), respectively. Consistent with depletion of the many actively replicating parasites from the 803 population by three daily sorbitol treatments, the 803 phasor distribution is shifted right with quiescence and increased levels of free NADH in persisters. The varied genetic backgrounds and baseline metabolic states of African GB4 and Southeast Asian 803 parasites help to explain different positions of the 803 and GB4 coordinates (p < 0.0001, Pillai trace).

The metabolic states of the DHA/sorbitol treated GB4 and 803 parasites were also assessed by FLIM-phasor analysis on Days 5 and 11 of dormancy (Fig. 4C). On these days, the mean coordinates of the GB4 and 803 distributions were located on the right side of the phasor plot. Although the coordinates of the mean phasor positions from each assessment can be affected by particulars of the culture conditions and experimental manipulations, we note that the large shift of the mean 803 signal to the right from its position at *t* = 50 hours to its position on Day 5 is consistent with the higher level of free NADH in the purer population of persisters (DHA *t* = 50 hours coordinates (0.63, 0.16) *vs*. DHA Day 5 coordinates (0.77, 0.21); p < 0.0001, Pillai Trace; compare Fig. 4B, right panel and 4C, left panel). The mean position of the DHA-treated 803 distribution then trended to the left, from its Day 5 coordinates of (0.77, 0.21) to Day 11 coordinates of (0.74, 0.22); p = 0.05, Pillai Trace; compare Fig. 4C, right panel and 4C, left panel). This trend suggests that a relatively greater level of bound NADH was developing on day 11, consistent with the presence of active ring forms waking from the 803 persister population.

## DISCUSSION

In the present study, we have used FACS sorting, ASM, and autofluorescence FLIM-phasor analysis to characterize signature changes in the mitochondria of *P. falciparum* persisters that develop after exposure to DHA. These mitochondria differ from the smooth and oblate mitochondria of actively replicating parasites by more rumpled and corrugated shapes that develop within 50 hours of DHA exposure. Along with their irregular shapes, the fluorescence volumes of persister mitochondria are often up to 5*-*fold larger and are also found in closer positions to the parasite nuclei than in active ring stages. Indeed, in many cases the ASM images show partial enwrapment of the nuclei by the mitochondria of the DHA-induced persisters. Previous studies on the ultrastructural effects of ART with various *Plasmodium* species *in vivo* found evidence for mitochondrial enlargement in electron microscopy images (32, 36-38). In other apicomplexans, mitochondrial enlargement has been reported in *Toxoplasma gondii* following ART pressure (33).

Is it possible that closer mitochondria-nuclei associations after DHA treatment could reflect an effect of diminished parasite volume and shrinkage of the 3D space available to the organelles? The larger erythrocyte surface areas of DHA-treated relative to untreated cells (Fig. S6) are not supportive of this possibility, although they cannot rule it out because the abilities of vehicle control- and DHA-treated erythrocytes to flatten might differ. Perhaps more important is that several key observations would remain unexplained by volume reduction that simply brings mitochondria closer to nuclei. The enlarged mitochondrial volumes and their rumpled and corrugated appearances would be unaccounted for by the physical effects of smaller containment. Cell volume reduction also would not explain the partial enwrapment of the nuclei by mitochondria, which in many cases extends across large surface areas of the nuclei.

Reactive oxygen species (ROS) capable of membrane damage are induced by ART in isolated *Plasmodium* and yeast mitochondria and this induction can be moderated by agents that reduce electron transport chain activity (39). Indeed, rapid killing is thought to result from depolarization of the parasite mitochondrial and plasma membranes upon exposure to ROS (40), and the mitochondrion has been proposed as a sensor of cellular damage produced by ART activity (41). Depolarization of mitochondrial membrane potential accompanies the disruption of mitochondrial function by ART in yeast (42). In human tumor cell lines, activation of the ART endoperoxide bridge by mitochondrial heme is thought to promote generation of ROS by the electron transport chain, leading to mitochondrial dysfunction and cellular apoptosis (43). Artemisinin-resistant *T. gondii* selected *in vitro* provided further evidence for the involvement of mitochondrial pathways, as demonstrated by amplifications of mitochondrial cytochrome b and cytochrome c oxidase I genes and mutations in the mitochondrial protease DegP2 ortholog (33). The same DegP2 ortholog was identified in a genetic screen: its disruption in *T. gondii* facilitated survival to lethal concentrations of DHA and its deletion in *P. falciparum* resulted in higher survival in the RSA (44).

Interestingly, in breast cancer cells and mouse embryonic fibroblasts the responses to mitochondrial stress include formation of tethers between nuclei and mitochondria, coined Nuclear Associated Mitochondria (NAM) (45). ROS production after exposure of THP-1 derived macrophages to the fungal toxin altertoxin II was followed by relocation of mitochondria to perinuclear regions (46). Similarly, we speculate that oxidative stress may be involved in triggering the development and survival of *P. falciparum* persisters through the morphological changes of mitochondria and their repositioning in closer association with nuclei.

Mitochondrial retrograde response (MRR) signaling provides communications from mitochondria to nuclei that promote cellular adaptations under conditions of stress (47). Contact microdomains between the two organelles involve cholesterol redistribution and may promote a survival response in breast cancer (45). The MRR pathway is conserved among species, as exemplified by nuclear protein CLK-1 in *Caenorhabditis elegans* and its homolog in human cells COQ7, which serves as a regulator of this pathway limiting ROS production (48). In these studies, mitochondria and the MRR were found to serve as barometers of cellular damage, capable of changing gene expression pathways to facilitate survival.

The changes in mitochondrial morphology, reduced mito-nuclear distances, and metabolic consequences induced by DHA exposure presented here may be tied to gene expression responses involved in persister formation as a survival mechanism. These changes appear to occur in PfK13 wild-type as well as C580Y mutants and may relate to an innate, multigenic growth bistability phenomenon whereby a fraction of the drug-exposed population switches into the metabolic quiescence of persister forms, as we have discussed elsewhere (19). How is it that less than ~1% (5, 7) of the parasite population becomes persisters while the larger fraction dies away? The answer may be a stochastic one that lies in systems of gene control. Feedback loops could rapidly flip the growth state of some cells to dormancy, producing forms that can survive DHA treatment while the actively replicating parasites succumb to the toxic effects of the drug. Understanding these feedback loops and the metabolic and functional features of persisters may lead to new chemotherapeutic approaches that can block clinical recrudescences.

## MATERIALS AND METHODS

### Materials and Methods

See **Text S1** in supplemental material.

## ACKNOWLEDGEMENTS

We thank Tom Moyer, Teresa Hawley and David Stephany in the Flow Cytometry Section of the Research Technologies Branch for their advice on FACS sorting, and Susan Pierce for supporting experiments at the Laboratory of Immunogenetics, NIAID. This work was funded by the NIAID Division of Intramural Research.

## COMPETING INTERESTS

No competing interests declared.

## SUPPLEMENTAL FIGURES

**FIG S1.**
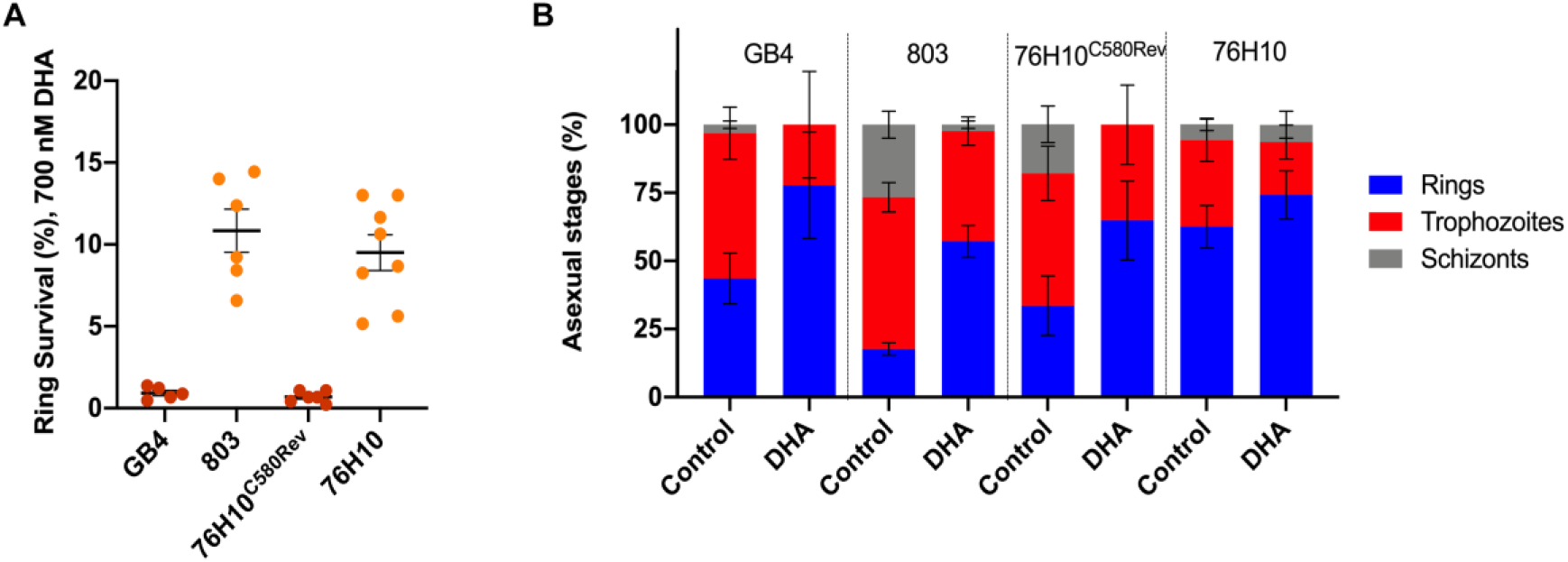
Independence of differential survival and delayed erythrocytic stage development after exposure of *P. falciparum* ring-stage parasite lines to 700 nM DHA for 6 hours. (**A**) Results of ring-stage survival assays (RSAs) performed on GB4, 803, 76H10^C580Rev^ and 76H10 lines. The GB4 line harbors a PfK13 K189T change upstream of a wild-type Kelch propeller region; 803 harbors the PfK13 C580Y propeller mutation; 76H10 is a progeny clone from GB4×803 cross that inherited PfK13 C580Y; and 76H10^C580Rev^ is an isogenic clone from 76H10 genetically edited to encode wild-type PfK13 C580. Zero-to-three hour ring-stage parasites were exposed for 6 hours to control vehicle (0.1% DMSO) or 700 nM DHA/0.1% DMSO treatment at *t* = 0 hours and percentage survivals were counted at *t* = 72 hours. At the *t* = 72 hour timepoint, the percentage survival was calculated for each strain by the ratio of surviving DHA-treated parasitemia to control vehicle-treated parasitemia. After DHA treatment, GB4 and 76H10^C580Rev^ parasites (C580 PfK13 propeller wild-type) survived at lower levels than the C580Y propeller mutant 803 and 76H10 parasites (0.93 +/-0.16 (mean +/-SEM, N=5) and 0.69 +/-0.14 (mean +/-SEM, N=6) *vs*. 10.84 +/-1.32 (mean +/-SEM, N=6) and 9.50 +/-1.09 (mean +/-SEM, N=8), respectively). (**B**) Stage distributions in control vehicle-treated or DHA-treated GB4, 803, 76H10^C580Rev^ and 76H10 parasite lines at *t* = 72 hours. The stage distributions were independently counted from Giemsa stained thin blood films by two observers blinded to the sample identifications. Results show a higher proportion of rings in DHA treated cultures relative to trophozoites and schizonts than in cultures treated with vehicle only. This higher proportion of ring-stage parasites in the second erythrocytic cycle suggests that the DHA treatment had temporarily slowed development of the PfK13 wild-type and C580Y mutant parasites for several hours in the first cycle. Such slowing is consistent with a previous study that found evidence for a delay in the *P. falciparum* erythrocytic cycle after ART exposure (Klonis et al., *Proc Natl Acad Sci U S A* **108**:11405-11410, 2011, https://doi.org/10.1073/pnas.1104063108). Transcriptomic data from the *P. falciparum* isolates of patients with acute malaria has likewise shown evidence for decelerated ring-stage development after ART treatment (Mok et al., *Science* **347**:431-435, 2015, doi:10.1126/science.1260403).

**FIG S2.**
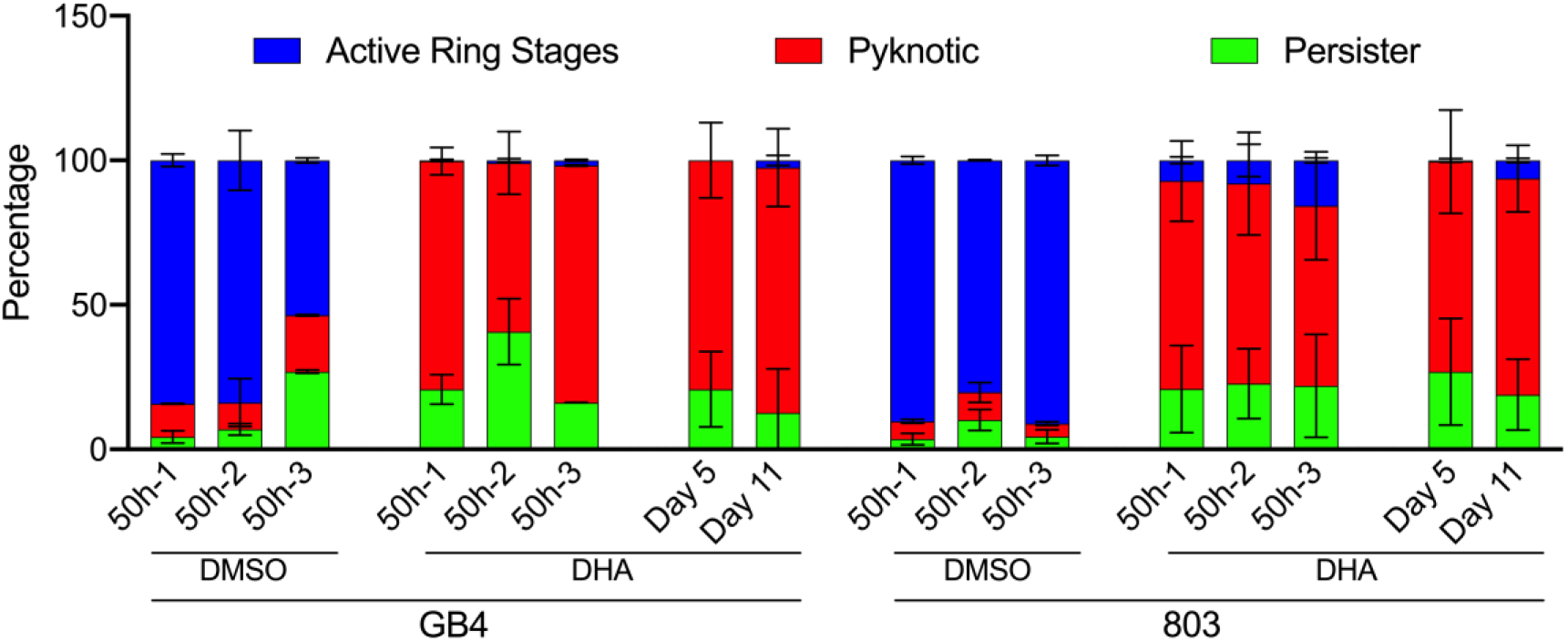
Proportions of parasites identified as active ring stages, pyknotic, and persister forms in *P. falciparum* 0.1% DMSO vehicle- and DHA-treated parasites at different collection timepoints. Counts were determined from Giemsa-stained thin blood films by two independent readers blinded to the slide identifications. The raw data counts were normalized to percentages from which the averages and standard deviations were calculated.

**FIG S3.**
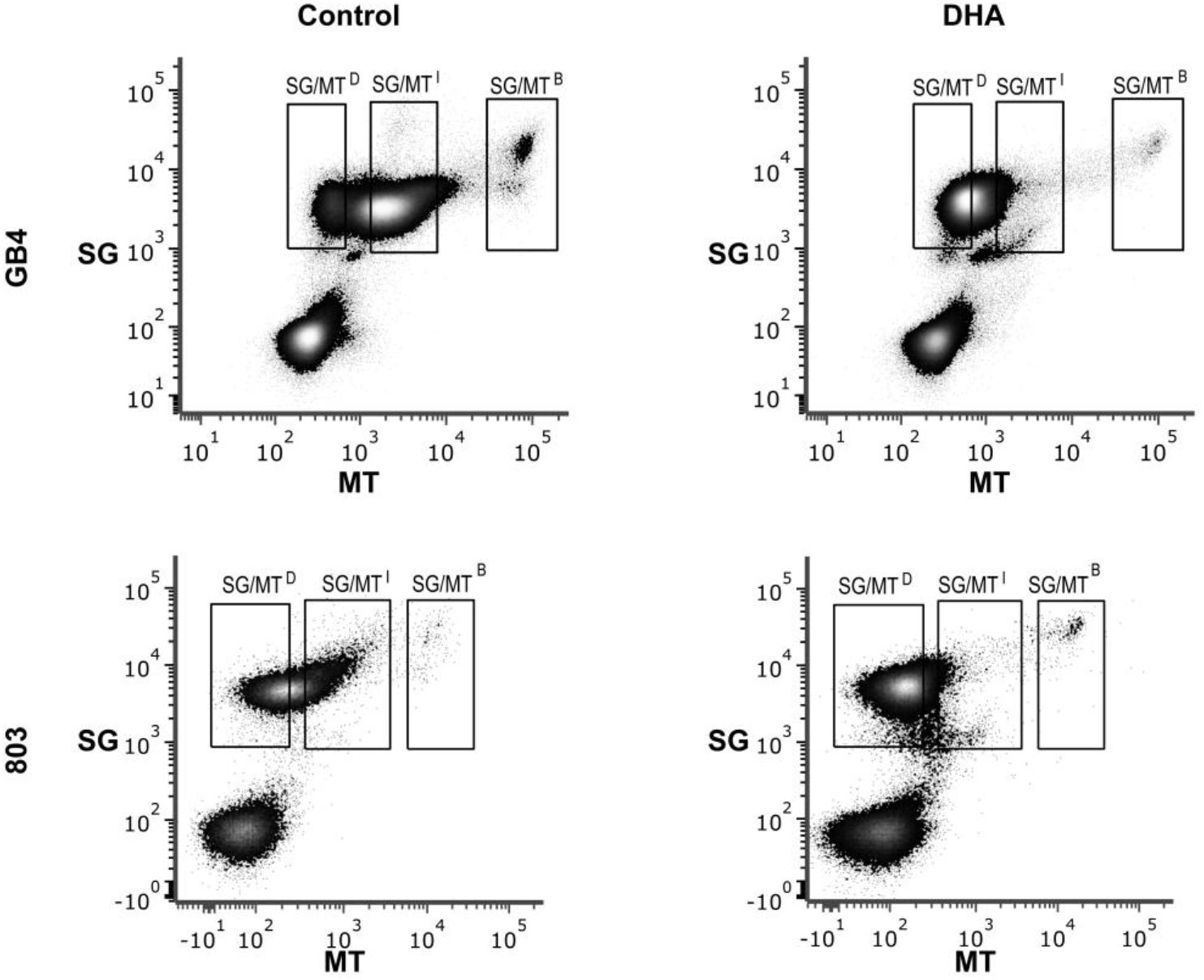
Gating strategy and distributions of the *P. falciparum*-infected erythrocytes collected for outgrowth experiments. Dot plots show the population groupings of control or DHA-treated GB4 and 803 parasites at experimental timepoint *t* = 50 hours. SG and MT axes indicate levels of SYBR Green I and MitoTracker Deep Red fluorescence, respectively. Gate labels identify the regions of dimly MT-fluorescent pyknotic cells (SG/MT^D^), ring-stage and persister parasites of intermediate fluorescence (SG/MT^I^), and brightly fluorescent late stages (SG/MT^B^).

**FIG S4.**
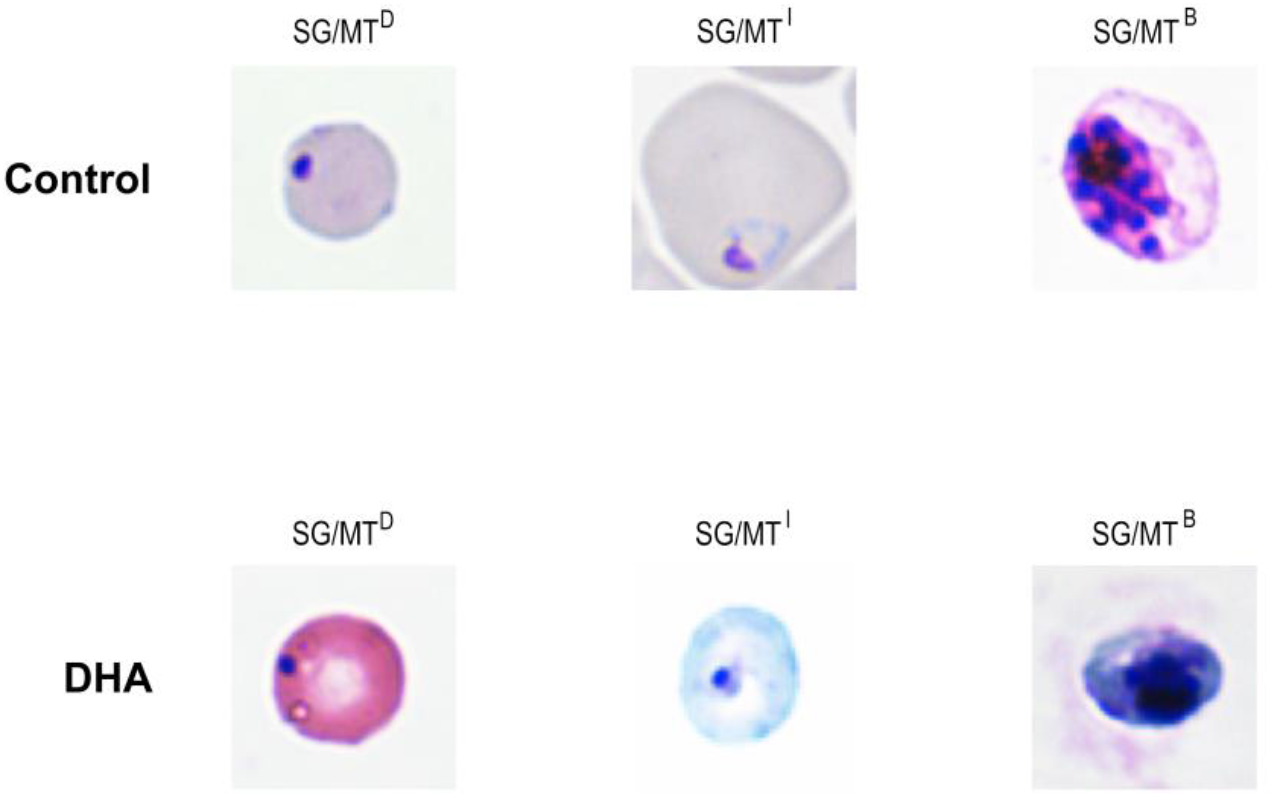
Representative images of GB4 infected RBCs after 0.1% DMSO vehicle or DHA exposure followed by FACS separation at *t* = 50 hours. The parasite-infected RBCS were washed twice with cRPMI before they were fixed in a thin blood film, Giemsa stained, and imaged by microscopy.

**FIG S5.**
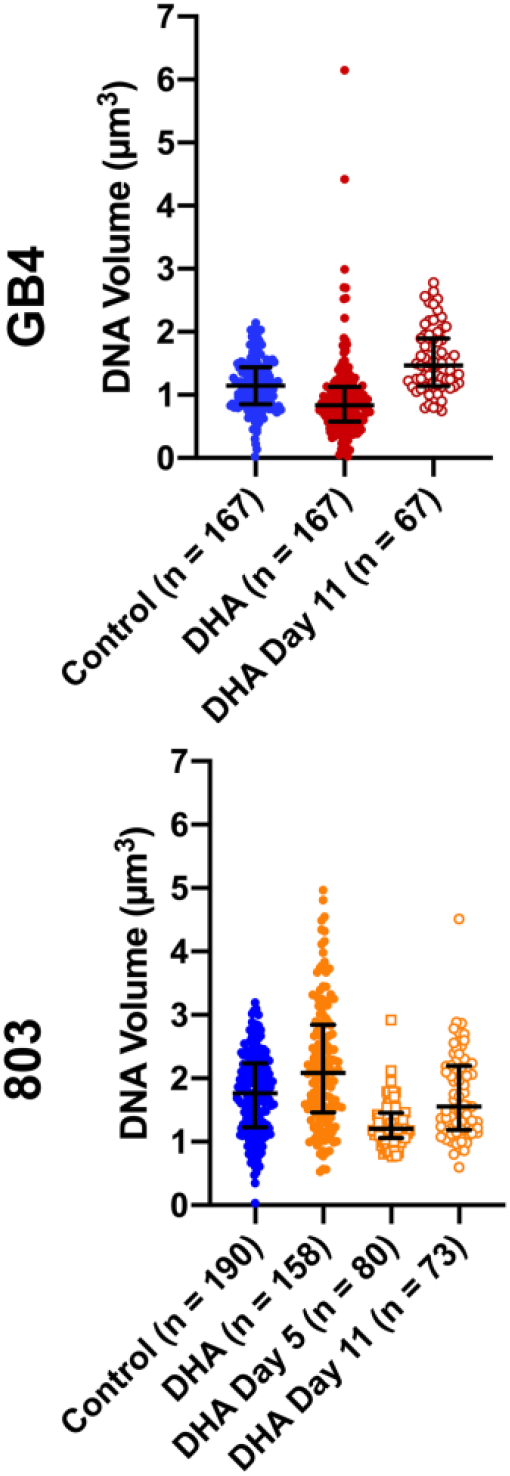
DNA volumes of 0.1% DMSO vehicle control or DHA-treated GB4 and 803 parasites at *t* = 50 hours, and of the DHA-treated parasites at Days 5 and 11 in the recrudescence studies.

**FIG S6.**
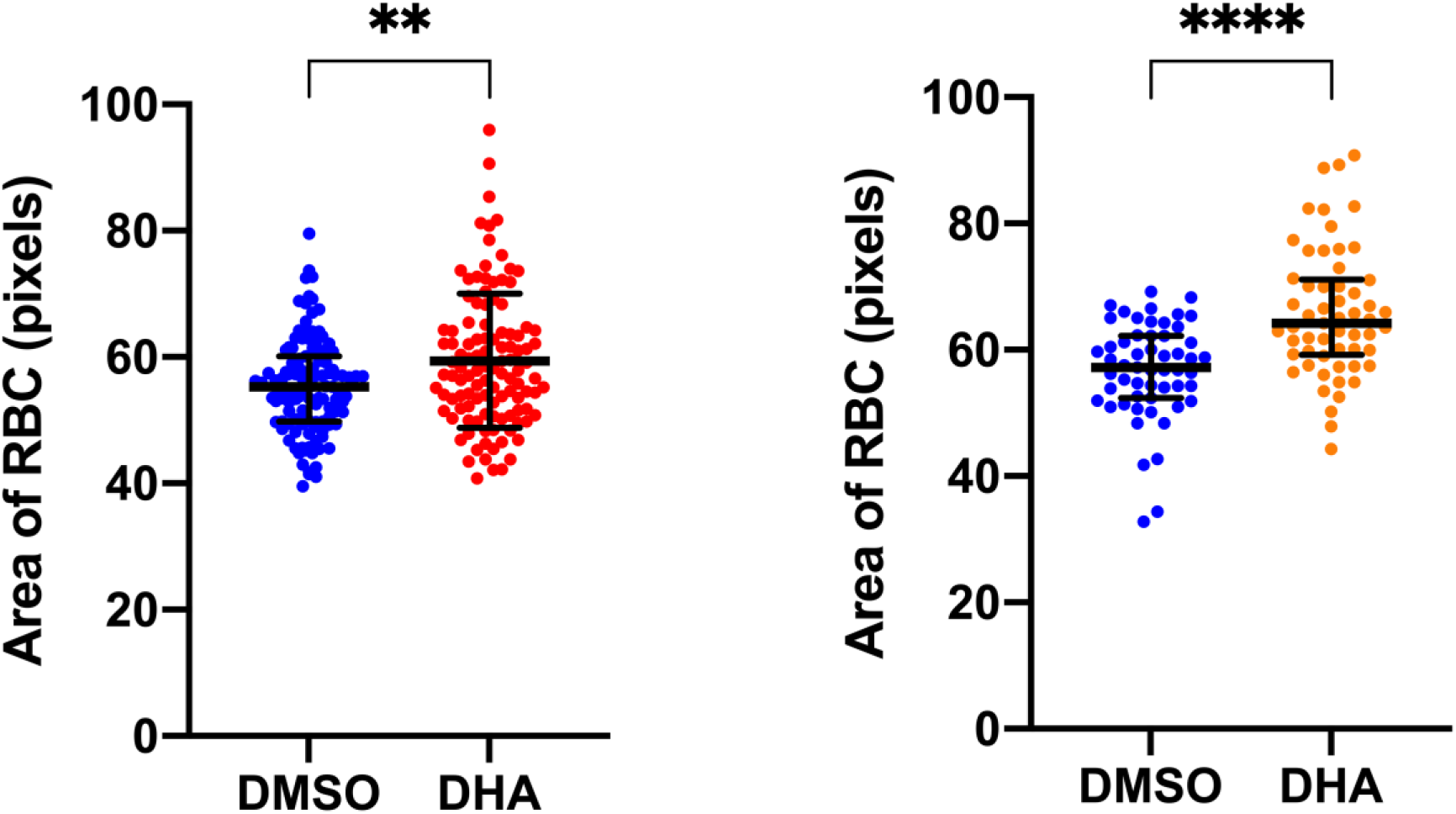
FACS sorted GB4 or 803 infected RBCs were seeded on a Poly-Lysine covered microwell. RBC areas were determined from DIC images of the infected erythrocytes at *t =* 50 hours after 0.1% DMSO vehicle control or 700 nM DHA exposure. Median plus IQR values were 55.28 (49.76-60.13, n = 115) and 57.84 (51.67-64.27, n = 113) for vehicle control- and DHA-treated GB4, respectively (p = 0.0012). Median plus IQR values were 57.18 (52.38-62.25, n = 55) and 64.25 (59.20-71.17, n = 57) for vehicle control- and DHA-treated 803, respectively (p < 0.0001)

## SUPPLEMENTAL TABLES

**TABLE S1.** Outgrowth of MT-fluorescent GB4 parasites after exposure to 700 nM DHA or 0.1% DMSO alone and sorting by FACS (replicate #1)

| Exposure | Fluorescence Intensity Group | Predominant Morphology | Seeded Cells/Well | Days to 0.5% parasitemia outgrowth |
| --- | --- | --- | --- | --- |
| 0.1% DMSO vehicle control | SG/MT<sup>D</sup> | Pyknotic Forms, some Ring Stages | 30,000 | 17 |
|  | SG/MT<sup>I</sup> | Ring Stages | 30,000 | 6 |
|  |  |  | 30,000 | 8 |
|  | SG/MT<sup>B</sup> | Trophozoites | 4,700 | 11 |
| DHA | SG/MT<sup>D</sup> | Pyknotic Forms | 30,000 | No Growth |
|  |  |  | 160,000 | No Growth |
|  | SG/MT<sup>I</sup> | Persister Forms and Ring Stages | 30,000 | 18 |
|  | SG/MT<sup>B</sup> | Trophozoites | 1,200 | 27 |

The vehicle-control or DHA treated GB4 parasites were sorted into groups by MT-fluorescence and collected in 1xHBSS/2% FBS. Cells were pelleted at 2500 rpm for 5 minutes, resuspended in culture medium, and seeded at the indicated numbers into 200 µl culture volumes (5% HCT) in the wells of a 96-well plate. Thin and thick Giemsa stained blood smears were monitored and assessed for parasitemia.

**TABLE S2.**
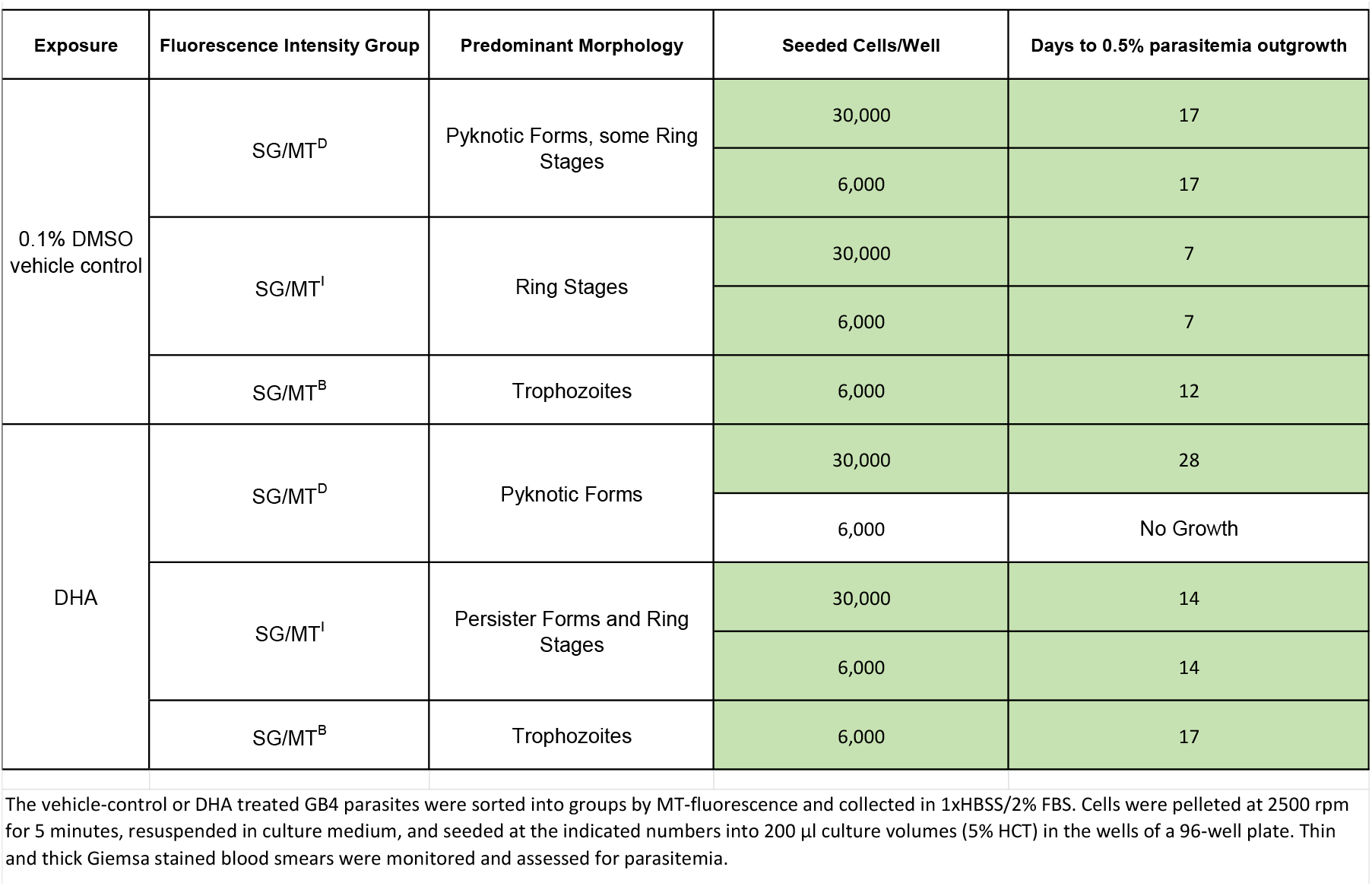
Outgrowth of MT-fluorescent GB4 parasites after exposure to 700 nM DHA or 0.1% DMSO alone and sorting by FACS (replicate #2)

| Exposure | Fluorescence Intensity Group | Predominant Morphology | Seeded Cells/Well | Days to 0.5% parasitemia outgrowth |
| --- | --- | --- | --- | --- |
| 0.1% DMSO vehicle control | SG/MT<sup>D</sup> | Pyknotic Forms, some Ring Stages | 30,000 | 17 |
|  |  |  | 6,000 | 17 |
|  | SG/MT<sup>I</sup> | Ring Stages | 30,000 | 7 |
|  |  |  | 6,000 | 7 |
|  | SG/MT<sup>B</sup> | Trophozoites | 6,000 | 12 |
| DHA | SG/MT<sup>D</sup> | Pyknotic Forms | 30,000 | 28 |
|  |  |  | 6,000 | No Growth |
|  | SG/MT<sup>I</sup> | Persister Forms and Ring Stages | 30,000 | 14 |
|  |  |  | 6,000 | 14 |
|  | SG/MT<sup>B</sup> | Trophozoites | 6,000 | 17 |

The vehicle-control or DHA treated GB4 parasites were sorted into groups by MT-fluorescence and collected in 1xHBSS/2% FBS. Cells were pelleted at 2500 rpm for 5 minutes, resuspended in culture medium, and seeded at the indicated numbers into 200 µl culture volumes (5% HCT) in the wells of a 96-well plate. Thin and thick Giemsa stained blood smears were monitored and assessed for parasitemia.

**TABLE S3.** Outgrowth of MT-fluorescent 803 parasites after exposure to 700 nM DHA or 0.1% DMSO alone and sorting by FACS.

| Exposure | Fluorescence Intensity Group | Predominant Morphology | Seeded Cells/Well | Days to 0.5% parasitemia outgrowth |
| --- | --- | --- | --- | --- |
| 0.1% DMSO<br>vehicle control | SG/MT<sup>D</sup> | Pyknotic Forms, some Ring Stages | 30,000 | 13 |
|  |  |  | 5,000 | 17 |
|  | SG/MT<sup>I</sup> | Ring Stages | 30,000 | 13 |
|  |  |  | 5,000 | 13 |
|  | SG/MT<sup>B</sup> | Trophozoites | 5,000 | 20 |
| DHA | SG/MT<sup>D</sup> | Pyknotic Forms | 30,000 | No Growth |
|  |  |  | 5,000 | No Growth |
|  | SG/MT<sup>I</sup> | Persister Forms and Ring Stages | 30,000 | 20 |
|  |  |  | 5,000 | No Growth |
|  | SG/MT<sup>B</sup> | Trophozoites | 5,000 | 22 |

The vehicle-control or DHA treated 803 parasites were sorted into groups by MT-fluorescence and collected in 1xHBSS/2% FBS. Cells were pelleted at 2500 rpm for 5 minutes, resuspended in culture medium, and seeded at the indicated numbers into 200 µl culture volumes (5% HCT) in the wells of a 96-well plate. Thin and thick Giemsa stained blood smears were monitored and assessed for parasitemia.

## Notes

### Competing Interest Statement

The authors have declared no competing interest.

### Summary of Updates

Text, figures, and supplementary information revised and updated.

