## Supplemental Methods for "Restructured mitochondrial-nuclear interaction in *Plasmodium falciparum* dormancy and persister survival after artemisinin exposure"

Running Title: **Mito-nuclear restructuring in *P. falciparum* persisters**

### MATERIALS AND METHODS

**Parasite Cultivation.** Human erythrocytes were obtained weekly from Virginia Blood Services (Richmond, VA). Blood was washed upon arrival with filtered RPMI 1640 media (containing 25 mM HEPES and 50 µg/mL hypoxanthine) (KD Medical, Columbia, MD), and stored at 50% hematocrit in a 4˚C refrigerator for use within a week from processing. *P. falciparum* GB4 (1) and 803 (2) parasites were grown in complete culture media (cRPMI) containing 1% Albumax II (Life Technologies, CA, USA), 0.21% sodium bicarbonate (KD Medical, Columbia, MD), and 20 µg/mL gentamicin (KD Medical, Columbia, MD) in RPMI-1640 (KD Medical, Columbia, MD), at 5% hematocrit, 37˚C, and a gas mixture containing 90% $N_2$, 5% $CO_2$, and 5% $O_2$. To monitor parasitemia, thin blood films were prepared on glass slides, fixed with methanol and stained for 15 minutes with 20% Giemsa staining (Sigma-Aldrich, St. Louis, MO). Using bright-field microscopy and a 100× oil objective, an estimated 1,000 erythrocytes were counted, and the number of parasitized cells was used to estimate percentage parasitemia. In recrudescence experiments, media was changed every other day and fresh erythrocytes were added every four days. When media was changed, blood films were made and used to monitor parasitemia.

**Ring-stage Survival Assay (RSA).** The 803 and GB4 cultures were tightly synchronized to obtained 0 – 3 hour ring stages through two successive 5% D-sorbitol treatments 46 hours apart (3). Each sorbitol treatment lasted 10 minutes at room temperature. Immediately following the second sorbitol treatment, cultures were adjusted to 2% parasitemia and 5% hematocrit in a total volume of 10 mL in a T25 flask (Thermo Fisher Scientific, Waltham, MA). Thin blood smears were prepared just before drug exposure. Cultures were then treated at a final concentration of 0.1% DMSO or 700 nM DHA/0.1% DMSO by addition of 10 µl of DMSO or 10µl of 700 µM DHA for 6 hours. After 6 hours of incubation, cells were washed and returned to culture in drug-free cRPMI. Sixty-six to ninety hours post DHA treatment, thin blood smears were prepared, fixed with 100% methanol, and stained with 20% Giemsa (Sigma-Aldrich, St. Louis, MO). Slides identifications were blinded from the investigators while parasitemia counts were obtained.

**Dihydroartemisinin Treatment and Magnetic Removal of Mature Stages.** The 803 and GB4 cultures were synchronized and treated with the control vehicle or DHA as above. Thirty hours after the initiation of treatment, the DHA treated culture was passed over a magnetically-activated cell sorting LD column (Miltenyi Biotec, Auburn, CA) to deplete the culture of parasites that escaped the DHA treatment and progressed past ring-stage. The same methodology as Teuscher et al. (4) was applied to the use of the LD column. Pelleted cells from culture were suspended in 2 mL of cold cRPMI and passed three times through the column over a period of 3 – 5 hours. Afterwards, the column was washed with 30 mL of cRPMI. Eluates were combined and pelleted at 2,500 rpm for 5 minutes, resuspended in 10 mL of cRPMI, and returned to culture. As the parasitized erythrocytes approached maturity, the cultures were placed on a rotator in the 37˚C incubator to minimize the occurrence of multiply-infected relative to singly-infected cells.

**Dihydroartemisinin Recrudescence Assay.** Tightly synchronized 803 and GB4 parasites, at approximately 2% ring-stage parasitemia, were treated with 700 nM of DHA for 6 hours. At 24, 48 and 72 hours post DHA treatment, the cells of both cultures were treated with 10 mL of 5% D-Sorbitol for 30 minutes at 37˚C and washed twice with cRPMI. This treatment removed mature stage parasites that survived the DHA treatment. Parasitemia was monitored by microscopy while media was changed on alternate days and fresh erythrocytes were added every four days until recrudescence.

**Fluorescence Associated Cell Sorting (FACS).** At $t$ = 50 hours, cells from 1 mL of each culture were pelleted and washed three times with 1ml of 1× HBSS (Gibco, Waltham, MA) containing 2% fetal bovine serum (FBS; Gibco, Waltham, MA). Washed samples were stained with 0.4× SG (Invitrogen, Waltham, MA) for DNA content and 0.1 µM MT (Invitrogen, Waltham,

MA) for mitochondrial potential (5). Samples for FLIM analysis were stained only with 0.1 µM MT to avoid SG spectral overlap (bleedthrough) with the endogenous FLIM signal (6). Control 0.1% DMSO vehicle-treated and DHA-treated samples were filtered through a 35 µm filter (Corning, Corning, NY) immediately before sorting.

Cells were sorted by using the BD FACS Aria Fusion, and gating was performed using the BD FACSDiva™ software (BD Biosciences, Franklin Lakes, NJ). For ASM, samples underwent two sorting steps: 1) isolation of infected from uninfected erythrocytes and 2) separation of parasite populations by DNA and mitochondrial signals. The first step enriched infected cells by collecting 1 million SG positive cells. Enrichment removed the vast majority of uninfected erythrocyte allowing for more accurate sorting of live and dead parasites in the second step. A highly pure sorting modality (4-way purity sorting for FACS Aria Fusion) was chosen for the second round of sorting. Enriched samples were collected in polystyrene round-bottom 5 mL tubes (Corning, Corning, NY) with 1 mL of 1× HBSS containing 2% FBS, pelleted at 2500 rpm for 5 minutes in the 5 mL polystyrene tube, resuspended, and sorted into two populations depending on the presence or absence of MT signal. Parasites in the PYK gate were used for a Giemsa-stained thin blood smear while parasites in the MT+ gate were used for ASM. All four samples were collected in polystyrene round-bottom 5 mL tubes (Corning, Corning, NY) with 1 mL of 1× HBSS/2% FBS.

For autofluorescence FLIM experiments, samples underwent a single step sorting. The MT+ cells were collected in polystyrene round-bottom 5 mL tubes (Corning, Corning, NY,) with 1 mL of 1× HBSS/2% FBS at room temperature and used immediately for imaging.

Data analyses were conducted and displayed using FCS Express (De Novo Software, Pasadena, CA).

**AiryScan Microscopy Analysis.** A 10µm x 10µm polydimethylsiloxane (PDMS) stencil microwell (Alvéole, Paris, France) was placed inside each well of a Lab-Tek 8 well chamber (Thermo Fisher Scientific, Waltham, MA). Ten µl of 0.01% poly-lysine (Sigma-Aldrich, St. Louis,

MO) was added to the microwell and incubated for 30 minutes at room temperature to allow poly-lysine to adhere to the slide. Afterwards, the microwell was washed twice with 10 µL of 1× HBSS to remove unbound poly-lysine. The DMSO and DHA-treated parasites positive for SG/MT were transferred to 1.5 mL Eppendorf tubes, spun down at 2,500 rpm for 3 minutes and resuspended in 20 µL of remaining wash buffer. Ten µL of cell suspension was added directly to the center of the poly-lysine coated microwell and incubated for 30 minutes at room temperature in the dark. Lastly, 200 µL of a 1:4 dilution of Intracellular (IC) Fixation Buffer (Invitrogen, Waltham, MA) in cell culture water (Gibco, Waltham, MA) was added to the chamber. The chamber was covered with a plastic membrane and stored at 4˚C for imaging within the following 24 hours. Parasites were imaged using the Zeiss LSM 880 with AiryScan. Afterwards, the ASM images were analyzed using Imaris 9.1.2 and volumes were adjusted. In the majority of cells, the small and low level mitochondrial DNA SG fluorescence was clearly distinct and excluded in the analysis. In a few cells, mitochondrial DNA fluorescence was not visible or was merged during thresholding of the DNA volume. Mitochondrial DNA regions were excluded if possible. The volumes of the mitochondrial and DNA volume were calculated as well as the distance between these volumes.

**Autofluorescence FLIM and Phasor Analysis.** When NADH undergoes multi-photon excitation, photons are released over time as NADH returns to its ground state, establishing a pattern of fluorescence decay that can be detected over a period of nanoseconds. The lifetime of decay differs for NADH in its free (0.4 nanoseconds) or protein bound state (1 nanosecond), which allows for the quantification of the redox state of the organism or tissue under study and display of the information in the phasor plane (7-9).

DHA-treated or control parasites stained with MT were seeded onto a microwell in an 8-well chamber coated with 0.01% poly-lysine. A confocal image was acquired using a Zeiss LSM 780 for each field of view with two channels: Differential Interference Contrast (DIC) and MT emission (633nm excitation and emission collected 640 – 740 nm range). Next, emission data were collected from the given field of view with the integrated Coherent Chameleon II Ti:Sapphire femtosecond-pulsed laser tuned to 750 nm. Photons for lifetime data fitting were collected using a Becker & Hickl TCSPC DCC-100 control card with two HCM-130 GaAsP hybrid detectors. A 525 nm dichroic separated the emission into the detectors and a 480/40 bandpass filter was used with the short wavelength detector. Each field of view was scanned until at least ten million photon events were collected. Phasor values were calculated for each pixel with the SPCImage software. The resulting files were exported, and a custom MATLAB algorithm used the signal from the MT confocal images to threshold the mitochondria, obtain the mean phasor values, and run statistical tests.

**Outgrowth Using Sorted Parasitized Cells.** Outgrowth experiments were used to investigate parasite viability after FACS. For this purpose, control and DHA-treated samples at $t$ = 50 hours were obtained through three gates: SG/MT Dim (SG/$MT^{D}$), SG/MT Intermediate (SG/$MT^{I}$) and SG/MT Bright (SG/$MT^{B}$) ( see Fig. S1 in the supplemental material). Each population was placed into a polystyrene tube with 1× HBSS/2% FBS at a final concentration of 1 mL. One hundred µL was taken from these samples for a blood film, and the cells in the samples were pelleted at 2,500 rpm for 5 minutes, resuspended in 180 µL of cRPMI, and transferred to a 96 well plate. Twenty µL of a 50% erythrocyte stock were added to each well to obtain a 5% hematocrit culture. Media were changed on alternate days and 150 µL of 3.3% hematocrit solution were added every 4 or 5 days to replenish erythrocytes lost to sampling. The parasitemia in each well was monitored through thick and thin smears, using 1 µL of parasitized erythrocytes for each smear, and the time it took for the parasitemia to reach 0.5% was recorded.

Parasite viability was also verified for both vehicle-treated control and DHA-treated GB4 and 803 parasites, utilizing the SG and MT stained samples.

**Statistical Analysis of ASM and FLIM-Phasor datasets.** Significance testing of the ASM datasets was performed by unpaired t-tests in Graphpad Prism version 8.4.3.

Autofluorescence FLIM-phasor datasets and mean phasor distribution coordinates were compared through Pillai trace calculations in MATLAB version 2019b.
